# Recording large-scale, cellular-resolution neuronal activity from freely-moving mice

**DOI:** 10.1101/2022.06.01.494442

**Authors:** Aniruddha Das, Sarah Holden, Julie Borovicka, Jacob Icardi, Davina Patel, Rushik Patel, Jacob Raber, Hod Dana

## Abstract

Current methods for recording large-scale neuronal activity from behaving mice with single-cell resolution require either fixing the mouse head under a microscope or attachment of a recording device to the animal’s skull. Both of these options significantly affect the animal behavior and hence also the recorded brain activity patterns. Here, we introduce a new method to acquire snapshots of single-cell cortical activity maps from freely-moving mice using a calcium sensor called CaMPARI. CaMPARI has a unique property of irreversibly changing its color from green to red inside active neurons when illuminated with 400nm light. We capitalize on this property to demonstrate cortex-wide activity recording without any head fixation or attachment of a miniaturized device to the mouse’s head. Multiple cortical regions were recorded while the mouse was performing a battery of behavioral and cognitive tests. We identified task-dependent activity patterns across motor and somatosensory cortices, with significant differences across sub-regions of the motor cortex. This new CaMPARI-based recording method expands the capabilities of recording neuronal activity from freely-moving and behaving mice under minimally-restrictive experimental conditions and provides large-scale volumetric data that are not accessible otherwise.

## Introduction

The mammalian brain processes sensory information using synchronized activity of brain-wide distributed neurons that are connected into local circuits^1-6^, which emphasizes the need for developing recording methods that are capable of capturing these complex activation patterns. To address this challenge, previous efforts have concentrated on improving the sensitivity of sensors to capture single-cell activity^7-11^ and enhancing the capability of recording systems to track neurons over large brain regions^12-16^. In parallel, scientific paradigms have shifted to analyzing neuronal activity in behaving animals while they process sensory cues to perform a task, with much of this work performed using the mouse as a model. This approach has enabled identifying the functional roles of specific cell types^17^, brain regions^4^, and/or projections between brain regions^1,6^, as well as determining how normal activity patterns are altered following a neurological condition^18^ or a neurodegenerative disease^19^.

Currently, recordings from many neurons spanning multiple brain regions with single-cell resolution in behaving mice are mostly conducted using genetically-encoded calcium indicators (GECIs). These experimental paradigms include either monitoring of head-fixed mice using two-photon laser scanning microscopy (TPLSM)^20,21^, or attaching a miniaturized imaging device to the skull of a freely-moving mouse to record single-photon or two-photon fluorescence^22-24^. Importantly, both of these methods have inherent limitations. For example, TPLSM recording from behaving rodents requires head fixation of the mouse under a microscope, which may result in activation of different neuronal circuits compared to natural, freely-moving behaviors^25^. In addition, state-of-the-art TPLSM systems are capable of recording from one large plane spanning up to several mm^2^, or from a few smaller, axially-shifted planes^14-16^. These microscopes are usually limited by the mechanics of laser scanning systems, which restrict the effective field-of-view (FOV) size that can be dynamically monitored. The acquired information is limited to the inside of this FOV, and so the activity of nearby neurons outside the FOV, even if they are labeled with a GECI, cannot be simultaneously detected. When brain activity is recorded using an implanted miniaturized imaging device, it allows head movement of the mouse during recording, but it puts a substantial weight on its skull. This additional weight may affect the mouse’s natural behaviors, and hence also the recorded neuronal activation patterns. In addition, the spatial resolution and volumetric recording capabilities are compromised compared to TPLSM recording.

Calcium-modulated photoactivatable ratiometric integrator (CaMPARI) is a calcium- and light-dependent fluorescent activity marker^26^, which may enable combining the relative simplicity of GECIs with TPLSM large-scale recording capabilities and free movement of the animal. Upon illumination with 400 nm light in a high-calcium environment, CaMPARI undergoes an irreversible conformational change, and its fluorescence emission changes from green to red in a process called photoconversion (PC). CaMPARI was previously demonstrated to label with red fluorescence active neurons in the mouse visual cortex based upon their tuning properties^26^, and a recent version of this sensor, CaMPARI2, exhibits a brighter signal and better contrast between the red and green components^27^.

CaMPARI’s unique calcium-dependent PC capability allowed us to generate a new experimental paradigm where the experimental recording and signal readout processes are separated. In this study, we show that this paradigm shift facilitates neuronal activity recording over a larger brain volume than what has been possible with state-of-the-art TPLSM systems. Large-scale CaMPARI-based recording is conducted by shining a PC light over the animal and its experimental environment, which induces red fluorescence in active neurons in the mouse brain in a non-transient manner. The readout of the photoconverted CaMPARI fluorescence is conducted after the experiment is completed using a standard TPLSM system. This recording paradigm is fundamentally different than previous recordings with TPLSM systems, where the recording and readout processes occur simultaneously and cannot be separated. In this study, we reveal the advantages of CaMPARI-based recording for detecting activity from brain volumes larger than 6 mm^3^ with single-cell resolution. We validate the accuracy of the CaMPARI-based recording by comparing the results to recordings with the widely-used GECI, jGCaMP7s^8^. We also demonstrate the capability of CaMPARI-based recording to monitor single-neuron activity over a large cortical volume in freely-moving mice without any mechanical device attached to the mouse during the recording phase, and to compare activity level patterns across five somatomotor cortical regions as the mice perform a battery of behavioral tasks.

## Results

### Characterization of CaMPARI-based recording capabilities

Although the green-to-red PC was reported to be permanent at the single-protein level^26,27^, we found that the photo-converted RGR *in vivo* decreased during the days following PC and ∼97% of it decayed by one week (Fig. 1A and Supp. Fig. 1), presumably due to degradation of the red protein and production of new (green) protein. To calculate the rate of the RGR decay, we longitudinally monitored V1 neurons (n=73 from 2 mice) and measured the RGR after PC. The results were fit with an exponential decay model with a half-life of 1.04 days (R^2^=0.99; Fig. 1B). Multiple PCs of the same brain region and neurons were demonstrated by two additional recordings, separated by 7 days each, and yielding similar activity patterns (Fig. 1C). Therefore, we concluded that CaMPARI can be used multiple times for sequential recording sessions.

**Figure 1:**
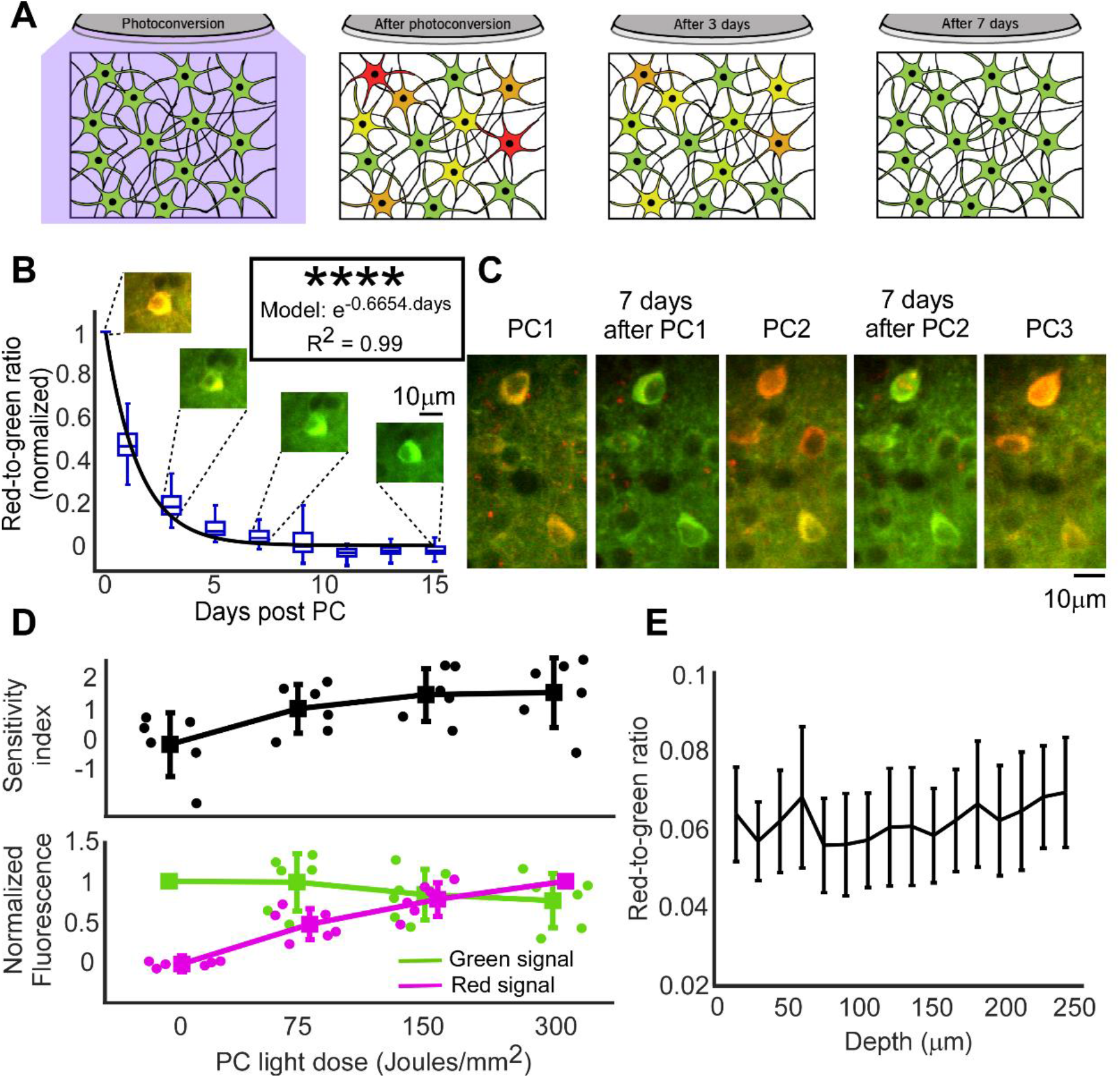
Characterization of CaMPARI-based recording. **(A)** Schematic illustration of the changes in CaMPARI green and red fluorescence during the week following photoconversion. **(B)** RGRs from photoconverted neurons (n=73 cell from 2 mice) were normalized to each cell’s RGR level immediately after PC (Day 0) and were monitored for 15 days. Median values were fit with an exponential curve (t_1/2_=1.04±0.076 days, mean±SE; R^2^=0.99; p=7*10^−10^, F-test vs. zero model). Example images of the same cell are presented above the respective days. **(C)** Example images of three repeated PCs in the same neurons. **(D)** Upper panel, sensitivity index (d’) values for separating V1 and S1 activity levels during visual stimulation have improved with increased PC light dose. Light dose experiments were performed sequentially, using the linearity of the PC process^26,27^, to calculate RGR for 75, 150, and 300 Joules/mm^2^. Circular dots show the median values from single recordings from all identified cells, and the large squares are the mean values across all single-experiment medians (data from n=3 mice, 24 PC sessions with 3-9 sessions per mouse; 207-762 neurons were recorded for each session with a median of 480 neurons/session). Lower panel, median green and red fluorescence signals from all recorded cells were normalized to their pre-PC levels (green) and 300 Joules/mm^2^ light dose (red) values. The green signal gradually decreased and the red signal increased with PC light dose (same data as in upper panel). **(E)** The median RGR across all recorded cells in each FOV, obtained from all mice and brain regions, showed no apparent differences across depths down to 240µm under the pia (n=5 mice, with data from 5 different motor and somatosensory regions, 2-110 cells/FOV, median=17). The solid line connects the average of single FOV median RGRs and error bars show the standard error of mean. No significant differences were found between different recording depths (one-way ANOVA, p=1.00).

Next, we characterized CaMPARI’s capability to identify active brain regions as a function of the amount of PC light shined on the mouse brain. Mice were injected with an AAV carrying the CaMPARI2 sequence into their primary visual and somatosensory cortices (V1 and S1, respectively; n=3 mice). The medians across the RGR recorded from all neurons in each region were compared after illumination of PC light during the presentation of a drifting grating movie to the contralateral eye (see Methods). Constant light quanta of ∼3.12 Joules/mm^2^ were delivered by illuminating a 7 mm diameter cross-section around the craniotomy opening using 120mW PC light for 1 sec, followed by 11 sec without illumination to allow the tissue to cool down. No signs of thermal damage were found for any tested light doses up to 300 Joules/mm^2^ (96 illumination cycles). The average V1 activity levels were higher than S1 for all tested light doses, and the sensitivity index (d’, see Methods) that quantifies the separation among the V1 and S1 RGR distributions improved with increased light dose (Fig. 1D, upper panel). As more PC light was used, the green CaMPARI fluorescence decreased down to 76% of its initial emission level and the red fluorescence increased (Fig. 1D, lower panel). Following these results, a light dose of 300 Joules/mm^2^ was used for the subsequent visual stimulation experiments to achieve optimal sensitivity. In a separate set of experiments, the RGR values were measured at different tissue depths down to 240 µm under the pia. No apparent changes in RGR values were found for different depths (Fig. 1E), which suggests that the PC-based recording levels were not biased by the tissue depth when layer II/III neurons were monitored.

### Simultaneous neuronal activity recording from multiple brain regions

Simultaneous volumetric recording over a large cortical area was accomplished by expressing CaMPARI2 in the monocular and binocular visual cortices (V1m and V1b, respectively) and the somatosensory cortices (S1) of the two hemispheres of mice (n=4). A drifting grating movie was presented to either the right or left eye of lightly-anesthetized mice and was synchronized with PC illumination that illuminated both hemispheres and covered a brain volume of 6 mm^3^ (∼10mm^2^ of cortical surface for each hemispheric window, with detectable signal down to a depth of ∼300 µm). Once the PC recording was completed, the neuronal RGR was recorded using TPLSM (Fig. 2A). A control group (n=2 mice) was expressed with jGCaMP7s in V1b, V1m, and S1 of one hemisphere and we recorded the fluorescence changes evoked by visual stimulation to each eye, as was previously done^7,8,28^. Both CaMPARI2- and jGCaMP7s-expressing neurons showed similar activity patterns, where visual regions were more active than somatosensory regions, and increased activity was detected in the contralateral compared to the ipsilateral visual regions (Fig. 2B-C, Supp. Fig.2-3). Notably, while CaMPARI-based data were recorded from all 6 brain regions simultaneously, jGCaMP7s-based data required a much longer recording process. Visual activity was recorded from 4-5 FOVs within each brain region sequentially, and therefore required combining 29 individual recordings.

**Figure 2:**
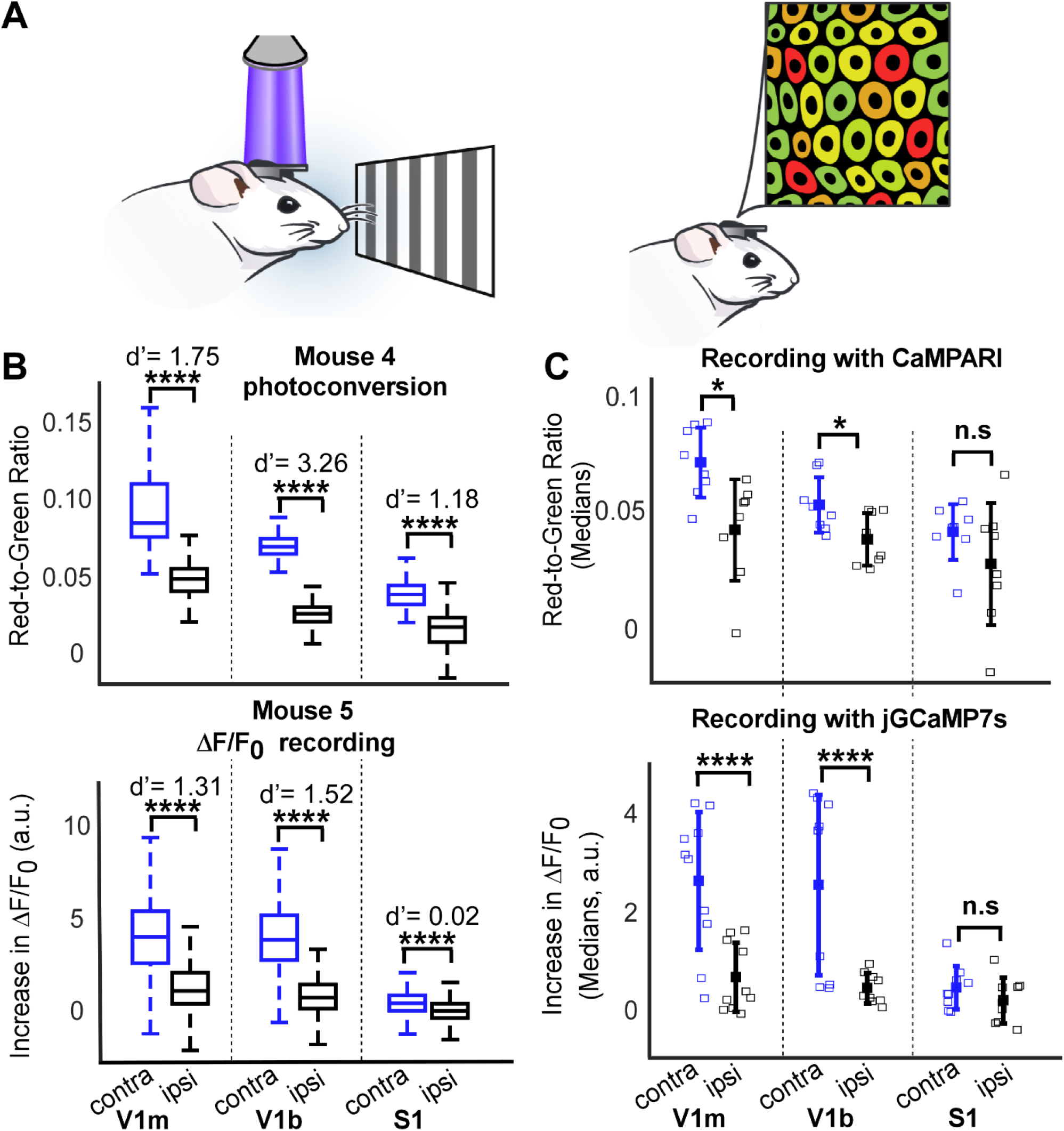
Simultaneous recording from multiple brain regions. **(A)** Schematic illustration of a visual-evoked activity recording experiment with CaMPARI. A drifting grating movie was presented to the mouse eye while PC light illuminated its two cortical hemispheres for recording neuronal activity (left). Following PC, the imprinted cellular RGR was read using TPLSM (right). **(B)** Example data from one mouse expressing CaMPARI2 (top) showing simultaneous recording data from V1m, V1b, and S1 in both hemispheres (111-256 cells/region, median=193). Activity levels are summarized by boxplots (horizontal lines show medians, boxes show the 25th–75th percentiles, whisker length is the shorter of 1.5 times the 25th–75th range or the extreme data point). Bottom, example data from a jGCaMP7s-expressing mouse showing similar distribution of increased fluorescence (930-2085 cells/region, median=1433). Sensitivity indices (d’) and significance levels (****, p<0.0001; Wilcoxon Ranksum test) across contralateral regions are plotted. **(C)** Summary of median activity levels measured with CaMPARI2 (top; n=4 mice, mice were recorded twice with stimulation for each eye; 81-1455 cells/regions, median=353; *, *p <0*.*05, paired t-test*). Recording from jGCaMP7s-expressing mice showed similar increases in fluorescence during stimulation (150-2085 cells/regions, median=634; see Methods).

### Large-scale volumetric recording of brain activity from freely-moving mice

Next, we moved to recording neuronal activity from freely-moving mice by expressing CaMPARI2 in 5 motor and somatosensory regions of the same hemisphere (motor caudal front limb, M_CFA_; motor rostral front limb, M_RFA_; motor neck-jaw, M_NJ_; somatosensory forelimbs, S_FL_; and somatosensory barrel field, S_BF_; n=6 mice; see Methods for details). The same mice were trained and tested on three tasks (novel object recognition in the open field (NOR), rotarod (RR), and fear conditioning (FC)), performing a new task every other week, using arenas equipped with a PC light source (Fig. 3A). For all experiments, the mice were first trained for the particular behavioral task. Following the completion of the training phase, the next session was conducted with the PC light turned on to photoconvert cells during 15 minutes of recording (Fig. 3B, Videos 1-3, see Methods for details). Readout sessions were conducted 24 hours after recording (Fig. 3C and Video 4). RGR was measured from all identified neurons from the pial surface down to a depth of ∼300µm. Evaluating cellular RGRs across different cortical regions and tasks in the same mouse (Fig. 3D), as well as across mice (Supp. Fig. 4), yielded a significant ∼2.5-fold increase in NOR median activity levels compared to RR and FC (Fig. 3E). Somatosensory regions were more active than motor regions across the three behavioral tasks (Fig. 3F). When comparing activity across the different motor regions, M_CFA_ was significantly more active than M_RFA_ and M_NJ_, although M_CFA_ and M_RFA_ project to the same limb^29^ (Fig. 3G). There were no significant differences in activity levels between the two somatosensory regions S_BF_ and S_FL_ (Supp. Fig. 6), or between cortical layers I and II/III. (Supp. Fig. 7).

**Figure 3:**
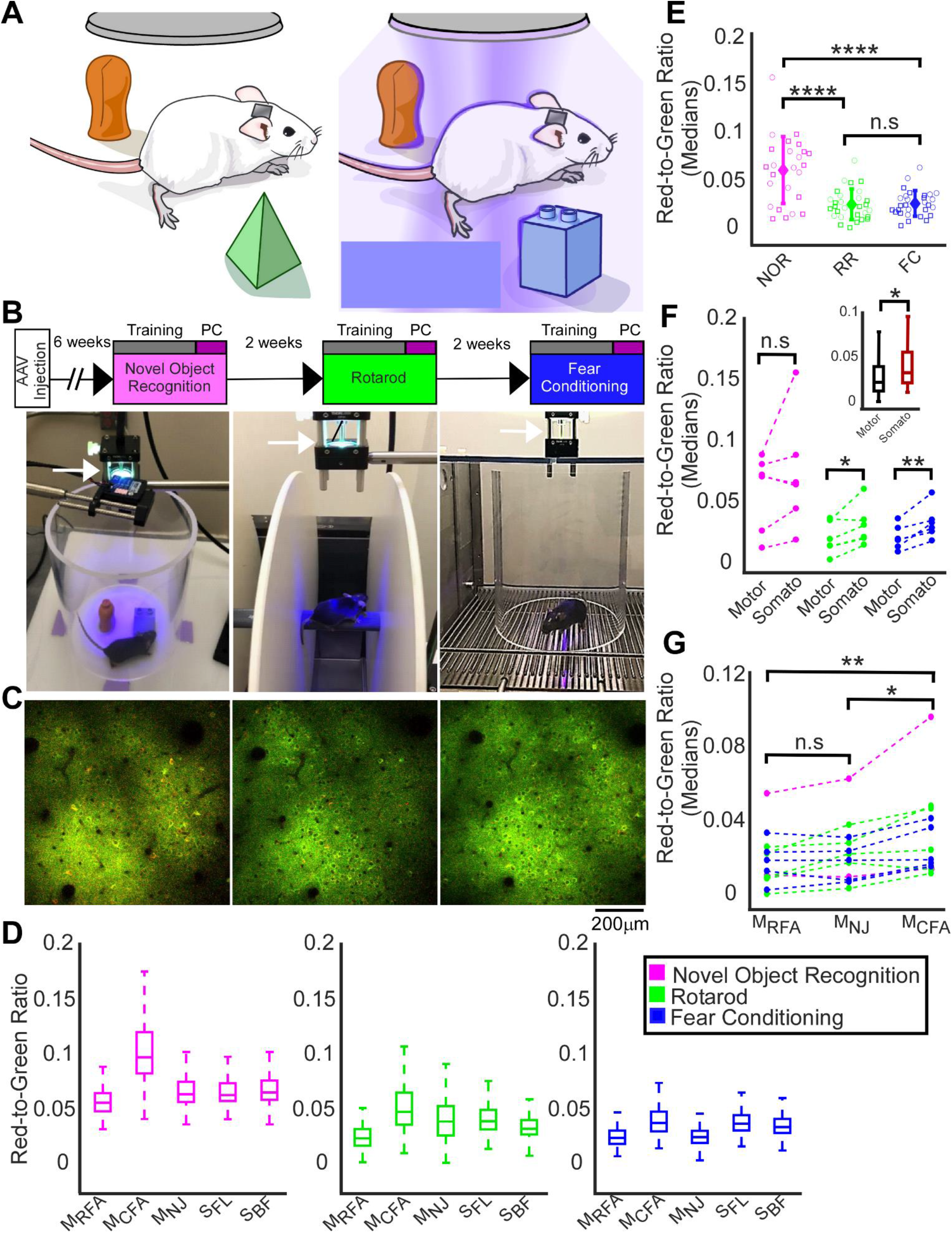
Recording large-scale, single-cell activity from freely-moving mice during behavioral and cognitive tasks. **(A)** Schematic illustration of the NOR experimental setup. A mouse trained on a task inside an arena without PC illumination (left). Following the completion of the training period, the PC light was turned on to record cortical activity while the mouse performed the task (right). RGR readout was conducted 24 hours after the experiment was completed in anesthetized mice using TPLSM. Each mouse was recorded every other week while performing three different tasks: NOR, RR, and FC. **(B)** Experimental timeline (top row) with images taken during NOR (left), RR (middle), and FC (right). The PC light source is indicated by white arrows. **(C)** Example post-PC images from the same brain region of the same mouse, which were acquired 24 hours after the recording of each task 14 days apart. **(D)** RGR from all identified cells in the five recorded cortical regions (same mouse as in **C**; NOR: 204-819 cells/region, median 547; RR: 166-847 cells/region, median 495; FC: 229-1315 cells/region, median 874). **(E)** Median activity levels (n=6 mice, five brain regions across motor and somatosensory cortical areas) during NOR were significantly higher than those during RR and FC (squares - motor, circles - somatosensory regions; mean ± s.d.; NOR: 42-906 cells/region, median 240; RR: 93-847 cells/region, median 251; FC: 85-1315 cells/region, median 321, **** *p < 0*.*0001*, Wilcoxon Ranksum test; n.s., non-significant). **(F)** Somatosensory regions were more active than motor regions across all tasks (inset). Across individual tasks, the increase was significant for RR and FC (n=6 mice; p =0.26 for NOR, p=0.036 for RR, and p=0.006 for FC; paired t-test, lines connect the pairs with data from the same mouse). **(G)** Activity levels in M_CFA_ were significantly higher than in M_NJ_ and M_RFA_ (p=0.018 and 0.006 respectively; n =5 mice, 89-902 cells/region, median 266; paired t-test with Bonferroni adjustment, lines connect the pairs with data from the same mouse). No significant difference in activity was observed between the M_NJ_ and M_RFA_ regions (Bonferroni adjusted p=0.16).

## Discussion

This work introduces a new method for acquiring simultaneous volumetric neuronal activity patterns from freely-moving rodents without mounting the animal’s head under a microscope or attaching a miniaturized imaging device to its skull. This new method enables large-scale recording across multiple brain regions at single-cell resolution. Therefore, this paradigm overcomes the respective limitations of TPLSM-based recording (optimized for planar data acquisition from head-mounted rodents) and miniaturized device-based recording methods (challenges with volumetric acquisition, image quality, and FOV size). This new method is especially beneficial for studying cortex-wide activity patterns in behaving mice, where minimal restrictions on the animal naturalistic performance is required. Importantly, CaMPARI-based recording is performed simultaneously across the entire PC-illuminated volume, and therefore may highlight brain regions or specific cell types that are active during specific behaviors. This large-scale mapping may serve as the first step to identify active brain regions (Fig. 2C and Fig. 3D-G) followed by a more detailed study of dynamic action potential firing patterns using either CaMPARI’s dynamic recording capabilities^26,27^ or by expressing a more sensitive GECI in the target brain region(s). CaMPARI-based recording is also compatible with mapping of cortex-wide activity patterns in response to different stimulations, including sensory, chemogenetic or optogenetic^30-33^.

Unlike existing recording methods, CaMPARI-based recording allows separation of the recording and readout processes. This separation allows conducting the recording by shining the PC light over an entire arena in which the rodent is contained, and to simultaneously record from all brain regions that are exposed to the PC light. Moreover, the return of RGR values to their baseline level within ∼7 days enables same-animal longitudinal monitoring, which differentiates CaMPARI-based recording from immediate early gene-based methods that require sacrificing the animal in order to read the activity data^34-36^. We note that future works may incorporate the use of the recently-published reversible CaMPARI (rsCamPARI)^37^, which will allow erasing the CaMPARI activity signal immediately after the readout session, and therefore eliminate the current restriction of CaMPARI2 with regard to the recording interval period.

The presented method is limited to acquiring snapshots of large-scale activity patterns. The CaMPARI2 sensor, which was used in this study, requires a relatively prolonged PC illumination time for achieving high-quality recording in freely-moving mice (up to 15 mins in this study). Shortening the recording session will further enhance the new method’s capability to monitor brain activity and highlight co-active brain regions across shorter time scales. Such improvement may be achieved by developing a new generation of the CaMPARI construct to address this specific challenge. In addition, in this study, we have expressed CaMPARI via intracranial AAV injection in specific cortical areas. Thus, we were limited to a finite number of brain areas that could be monitored. The cortex-wide recording capabilities of CaMPARI could be maximized by using systemic AAV injection^38^ or by developing a transgenic CaMPARI mouse line. Both of these options will enable expressing CaMPARI over most of the mouse cortex, which would achieve access to approximately 1 million neurons without changing the presented recording protocol^39^.

Finally, we established recording from the same mice performing three different behavioral and cognitive tasks, and show changes in brain activity patterns across tasks and brain regions, with similar activity levels in recorded cortical layers (Fig. 3). This type of data demonstrates that CaMPARI-based recording facilitates longitudinal *in vivo* neuronal activity studies during non-restricted behaviors in the same animals. With the recent increase in interest in studying large-scale brain activity patterns, and specifically the characteristics of distributed neuronal circuits^40-42^, the presented method adds unique capabilities and complements the tools that are available to the neuroscience community.

## Supporting information

Das et al. supplementary information

## Author contribution statements

HD and JR conceived the project. HD, JR, and AD designed the study, and interpreted the results. AD performed all surgeries, characterization, and visual stimulation experiments. SH recorded from freely-moving animals. AD, JB, JI, DP and RP analyzed the data. AD, HD, and JR wrote the paper with comments from all authors.

## Acknowledgements

The authors would like to thank Drs. Eric Schreiter and Benjamien Moeyaert and the HHMI Janelia GENIE project for developing and sharing the CaMPARI sensor. We thank Dr. Stefanie Kaech Petrie for her assistance with conducting the TPLSM recording experiments. We thank Drs. Christopher Nelson, Dimitrios Davalos, and Shy Shoham for helpful comments and suggestion to improve the manuscript. JR and HD are supported by grants R21AG065914 and U01NS123658 from the NIH.

## Methods

All experimental and surgical procedures were performed following the set guidelines and protocols approved by the Lerner Research Institute and Oregon Health & Science University Institutional Animal Care and Use Committees (IACUCs) and Institutional Biosafety Committees (IBCs) and were consistent with the ARRIVE guidelines. Mice were group-housed in standard vivarium conditions until the start of the study. The vivarium was maintained at 20-21°C and food (in LRI facility: Teklad 2918 regular diet, Envigo; in OHSU facility: PicoLab Rodent Diet 20, no. 5053; PMI Nutrition International) and water were available *ad libitum*. Lights were kept on a 12-hour light/12-hour dark cycle, and experiments were conducted during the light time.

### Surgical procedure and virus injection

For recording visual-evoked activity (Fig. 2), 8-10-week-old C57BL6/J mice (n=4 females) were anesthetized using isoflurane (3% for induction, 1.5% during the surgery) and placed on a heating pad. Each mouse was injected with local pain medication (bupivacaine 0.5%) and the skull bone above the two cortical hemispheres was exposed. A 3×5 mm^2^ craniotomy was drilled (Omnidrill35, World Precision Instruments) over an area covering the monocular and binocular primary visual (V1m and V1b, respectively) and primary somatosensory cortices (S1) in both hemispheres. An adeno-associated virus (AAV) solution expressing the CaMPARI2 sensor (*hsyn*-NES-his-CaMPARI2-WPRE-SV40, Addgene catalog number 101060) was injected into two locations, separated by ∼600µm, in each cortical region (50 nL of ∼1× 10^12^ GC/mL solution, 3 injection depths per location, 200µm, 400µm, and 600µm under the pia) using an automated injection pump (Fusion 200 touch Syringe Pump, Chemyx, and Micro-2T, WPI) and a pulled and beveled micropipette (P-1000 and BV-10, respectively, Sutter Instruments). Injection coordinates were chosen according to the mouse brain atlas^43^: 2.2 mm lateral and 0.2 mm anterior to Lambda (V1m), 2.8 lateral and 0.2 mm anterior to Lambda (V1b), and 2.5 mm lateral and 3.4 anterior to Lambda (S1). Cortex buffer^44^ was used consistently to keep the brain wet during the time of surgery and injections. Following the viral injection, a cranial window (two glued layers of rectangular glass, Tower Optical Corporation) was placed carefully, and a custom-made metallic head bar was attached using dental cement (Contemporary Ortho-Jet, Lang Dental). Animals were injected with Buprenorphine (0.1mg/kg) and Ketoprofen (5mg/kg, immediately, 24, and 48 hours after the surgery) for post-operative care and were allowed a minimal recovery time of 3 weeks before the start of experiments.

For recording activity from awake, behaving mice (Fig. 3), a similar surgical procedure was used (n=6 mice, 4 males and 2 females), but the AAV was injected into 5 locations into the mouse left motor and somatosensory cortices, according to microstimulation study^29^ and mouse brain atlas coordinates^43^: 1) 1.25/2 mm lateral/anterior to Bregma (rostral forelimb area of the motor cortex, M_RFA_); 2) 1.25/0 mm (caudal forelimb area of the motor cortex, M_CFA_); 3) 2/1.5 mm (neck and jaw regions of the motor cortex, M_NJ_); 4) 2/0 mm (forelimb region of the primary somatosensory cortex, S_FL_); and 5) 2.5/-1.75 mm (somatosensory barrel field cortex, close to the border with the trunk region, S_BF_). AAV solution (40 nL) was injected at depths of 250µm, 500µm, and 750µm in each location.

### Recording of visual-evoked activity

Mice were lightly anesthetized (0.5% isoflurane), held on a 37°C heating pad, and injected with Chlorprothixene Hydrochloride (IM, 30 µl of 0.33 mg/ml solution, Santa Cruz). PC of the CaMPARI2 signal started at least 30 min after the Chlorprothixene Hydrochloride injection, and after verifying that the mouse was responsive to pain but not voluntarily moving. The visual stimulation was presented to the mouse’s right or left eye and generated using the psychophysical toolbox^45,46^ in Matlab (Mathworks) on an LCD monitor (30×36 cm^2^ display, located 15 cm in from of the mouse right eye, tilted 45° with respect to the nose line, and covered with a blue plexiglass to minimize contamination into the recording channels) that subtended an angle of ±50° horizontally and ±45° vertically. The visual stimulus consisted of a drifting grating moving in 1 of 8 directions for 4 sec, followed by 8 sec of gray display. This stimulation cycle was repeated 5 times. PC light was delivered for 1 sec during the presentation of the drifting grating, 1.5 sec after the grating appeared, using an X-Cite Fire lamp (Excelitas) and a 40/40 nm bandpass filter (Brightline, Semrock) with 120 mW output at the sample plane. For recording of visual-evoked activity, the PC light covered a ∼12 mm diameter circle that included the two cranial windows in it, with an intensity of ∼0.9 mW/mm^2^. After PC was completed, CaMPARI2 signal was recorded using a two-photon microscope with resonant/galvo scanners (Bergamo II, Thorlabs) with 1040 nm excitation light (Insight X3, Spectra-Physics). Images were acquired using ThorImage 4.1 software (Thorlabs) with 15 frames per second and 1024×1024 pixels covering an area of 585×585 μm^2^ of layer II/III neurons. Green and red CaMPARI2 signals were recorded simultaneously (525/50 nm and 607/70 nm filters, respectively, separated by a 562 nm dichroic filter, Semrock) using 2 GaAsP PMTs (PMT2100, Thorlabs). For measuring the decay of red CaMPARI signal over time, the same V1 neurons were monitored immediately after the PC and over the subsequent 15 days.

For measuring visual-evoked fluorescence changes with jGCaMP7s^8^, the same TPLSM system described above was used with acquisition of 30 frames per sec, 512×512 pixels, and the same FOV size. The same drifting grating movie (with 4 sec of drifting grating followed by 4 sec of gray display) was presented to either the right or the left eye of the mice, and activity was measured from V1b, V1m, and S1 regions of the left cortical hemisphere to sequentially acquire ipsilateral and contralateral activity data.

### Recording of cellular activity from freely-moving animals

Mice were implanted with one cranial window over their left hemisphere as described above. Following the craniotomy, mice were given 7 days for recovery and were shipped from the Lerner Research Institute to the Oregon Health and Science University, where they were given an additional 3 weeks for quarantine and recovery. Mice were then randomly divided into 2 groups. Each week, one group was trained for 2-3 days on one of three behavioral tasks (see below). Then, the mouse cranial window was illuminated with PC light during the last trial of a given behavioral test. A broadband light source (XCite Xylis, Excelitas) with a 400/40nm bandpass filter (FBH400-40, Thorlabs) and lightly focused by a 100 mm achromat lens (AC254-100-A, Thorlabs) was placed 20 cm above the arena to illuminate a circular cross-section of 15.25 cm in diameter in which the CaMPARI PC occurred. The light intensity was 485 mW, or ∼2.65 mW/cm^2^. Mice were placed inside a plastic enclosure (16.5 cm diameter; TAP plastics) on matte white plastic flooring to keep them inside the illuminated region, and 15 min of PC illumination were used for each recording to elicit sufficient PC signal (based upon preliminary experiments we conducted with other mice to calculate the required light dose and duration).

Activity readouts were acquired using two-photon microscopy 24 hours after the PC during the behavioral and cognitive tests. Mice were anesthetized with isoflurane (4% induction, 1.5% maintenance) and put on a heating pad with core body temperature monitored by a rectal thermometer. Imaging was conducted with a Zeiss LSM 7 multiphoton microscope (Zeiss instruments), which utilizes a femtosecond-pulsed Ti:Sapphire laser (Chameleon Ultra II, Coherent), two BiG.2 GaAsP detectors, and Zen imaging software (Zeiss). The laser was tuned to 1040 nm excitation wavelength at 16mW when recording from the brain surface and up to 24mW when imaging 300 µm under the pia. Distilled water was placed on the cranial window to image with a 20x/1.0 water immersion objective (Zeiss, 421452-9880). We acquired volumetric data of layer I and II/III neurons (z-stack of ∼100 images typically from the brain surface, 512 × 512 pixels, 425 µm FOV size, 3µm step size between adjacent images) with green and red channels (500-550nm and 575-610nm, respectively, with a 560nm dichroic filter) from all identified CaMPARI2-injected regions in all animals.

### Behavioral testing

**1)** Exploratory behavior, measures of anxiety, and object recognition were assessed in an open field for a total of 4 consecutive days. Days 1 and 2 included exposure to the open field without objects for 5 min/day. Introduction of two identical objects occurred on day 3, and day 4 consisted of replacing one of the familiar objects with a novel one and 400nm illumination over the entire arena. Both days with objects (3-4) consisted of 15-minute trials. Behavior was recorded with Ethovision 15 XT software. Camera and PC light-guide suspensions over the arena were built with Thorlabs mechanical component.

**2)** Sensorimotor performance on the rotarod was tested for three consecutive days containing three trials each. Days 1 and 2 consisted of a standard rotarod protocol, with a starting speed of 5 RPM that was increased by 1 RPM every 3 seconds. On the third day, parameters were adjusted to a starting speed of 4 RPM, a maximum speed of 12 RPM, and an increase of 1 RPM every 60 seconds. This ensured that the animals remained on the rod for 5 minutes per trial (3x trials per day) and allowed for full 15-minute illumination while on the rod. Experimenters noted the speed at which the rod was turning when the animal fell off and the duration of time that the animal was able to stay on the rod before falling.

**3)** Contextual fear memory over two days was tested using a Med Associates mouse fear conditioning system for use with optogenetics (MED-VFC-OPTO-USB-M, Med Associates). During the training day, the animal was placed in the plastic arena (described above) that was located inside a white LED-lit (100 lux) fear conditioning chamber with a metal grid floor. Animals were habituated to the arena for a 300-s baseline period, followed by a 2-s, 0.7mA foot shock, administered a total of 4 times at 60-s intervals. The following day, animals were placed in the same plastic enclosure within the fear conditioning chamber and contextual fear memory recall was tested for 15 minutes under illumination. Movement in the chamber and the percentage of freezing were automatically determined by the Med Associates VideoFreeze automated scoring system (MED-SYST-VFC-USB2, Med Associates).

### Data analysis

Data analysis was performed using custom MATLAB scripts. For visual stimulation experiments and depending upon the specific experiment, cell bodies were segmented for each FOV using either a semi-automatic algorithm^7^, Suite2P softeware^47^, or the CellPose software^48^. Both of the two latter options allow for automatic segmentation with experimenter quality control. Suite2P was used for segmenting experiments where dynamic recording with jGCaMP7s was used, Cellpose was used to segment data from PC experiments with CaMPARI2, and semi-automatic segmentation was used when one of the automatic approaches yielded poor results. We compared the performance of the three segmentation methods on identical data and found that they produced similar results except for the number of detected cells, which was higher for Suite2P (Supp. Fig. 8). For each recording day, we calculated and corrected the effect of dark current values for the green and red PMT channels by recording images for each channel with the same exposure parameters used for the experiments, but with the laser turned off. Since CaMPARI’s green signal penetrates the red channel^26^, we calculated and corrected for the red-to-green contamination ratio by recording CaMPARI images before PC (pre-PC) and measuring the contamination ratio. Green-to-red corrections values were 13.8% and 6.2% for the data recorded in the Lerner Research Institute and Oregon Health and Science University, respectively. Finally, since the angle of the cortical window on each hemisphere was different with respect to the microscope’s optical axis, we measured a weak component of red channel auto-fluorescence for each hemisphere that was corrected by finding the intercept of the linear regression line of the green and red signals from cells from the same hemisphere before PC (Supp. Fig. 9). Following these corrections, we calculated the post-PC red-to-green ratio (RGR) for all cells in a brain region and used the median value for comparisons across regions. We calculated the sensitivity index (d’) to measure the separation between visual and somatosensory cortical regions following the formula mentioned below:

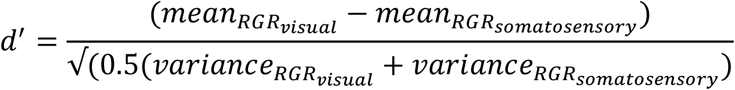

For light dose calculations, we consecutively photoconverted the same brain region several times on the same day and measured the RGR from all cells after each PC. The light dose was calculated as the total light intensity for each PC event divided by the illumination cross-section and multiplied by the illumination time.

For calculating visual-evoked fluorescence changes measured by jGCaMP7s, we integrated the fluorescence changes (F_resp_) above the baseline level (F_base_) during all appearances of the drifting grating stimulation for each recorded neuron by using the formula [sum(F_resp_ /2) – sum(F_base_)]. F_resp_ was measured over the 4 sec where the visual stimuli were presented, and F_base_ was measured over the 2 sec before the appearance of the respective drifting grating.

For recording from freely-moving mice, we first averaged every 3 adjacent images (spanning 6 µm in the z-axis) and skipped the next 2 images, in order to generate a set of images with minimal overlap of cells. We performed the same signal corrections as described above and calculated the RGR for all recorded cells and the median values for comparing across brain regions, mice, and tasks.

