## Supplementary material for "Recording large-scale, cellular-resolution neuronal activity from freely-moving mice": Das et al. supplementary information

### Supplementary Figures

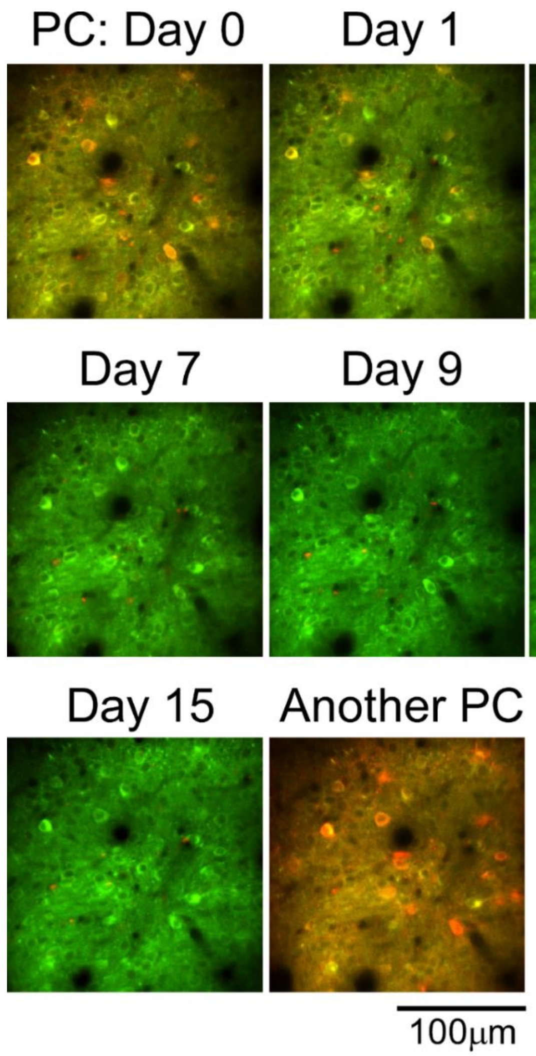


**Supplementary Figure 1. Decay of RGR over time.** Images of the same FOV were captured immediately after PC during visual stimulation (Day 0) and over the following 15 days. After the last recording session on Day 15, CaMPARI was again photoconverted using the same visual stimulation ("Another PC").

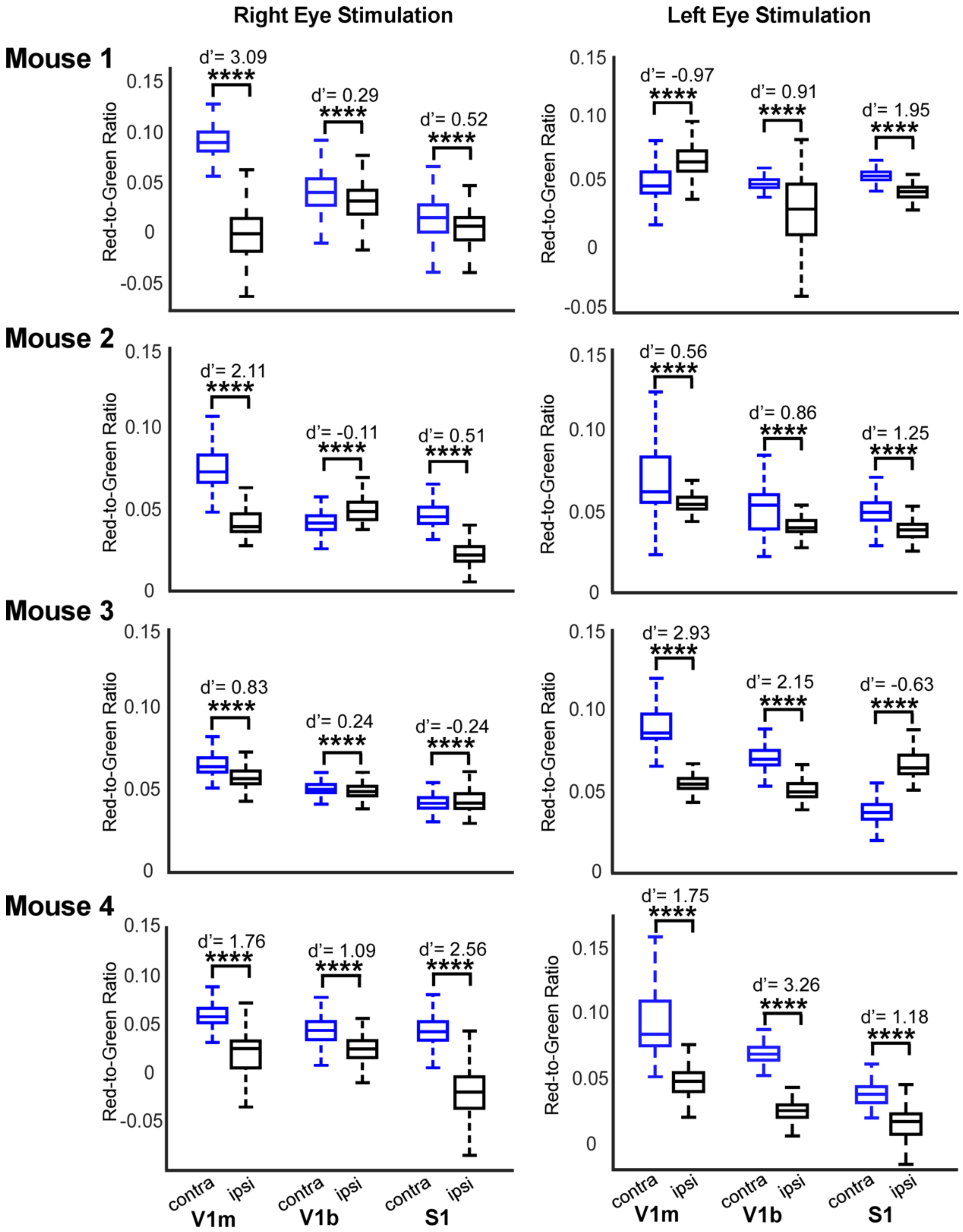

Right Eye Stimulation
Left Eye Stimulation
Mouse 1
Mouse 2
Mouse 3
Mouse 4
Red-to-Green Ratio
d'= 3.09
d'= 0.29
d'= 0.52
d'= -0.97
d'= 0.91
d'= 1.95
d'= 2.11
d'= -0.11
d'= 0.51
d'= 0.56
d'= 0.86
d'= 1.25
d'= 0.83
d'= 0.24
d'= -0.24
d'= 2.93
d'= 2.15
d'= -0.63
d'= 1.76
d'= 1.09
d'= 2.56
d'= 1.75
d'= 3.26
d'= 1.18
****
contra
ipsi
V1m
V1b
S1

**Supplementary Figure 2: Single-mouse data of visual-evoked responses in each brain region show increased activity in contralateral V1 regions, as recorded with CaMPARI2.** Summary of RGR for n=4 recorded mice. Each mouse was recorded twice for detecting activity patterns over 6 brain regions in 2 hemispheres after stimulation of either the right or the left eye. For most recordings, activity of contralateral V1m and V1b regions was higher than their ipsilateral counterparts, with significant increases in the median activity level across all recordings. Changes in S1 activity were significant for single recordings, but not when comparing all recordings. Data from mouse 4 left eye is also presented in Fig. 1D, and the summary of median activity levels from all mice is shown in Fig. 1E of the manuscript. Values of the sensitivity index (d') and significance (Wilcoxon Ranksum Test) of comparison across the same brain regions in the contralateral hemispheres are shown (Mouse 1: 107-1219 cells/region, median= 429; Mouse 2: 81-456 cells/region, median= 324; Mouse 3: 214-1455 cells/region, median=868; Mouse 4: 111-350 cells/region; median=249).

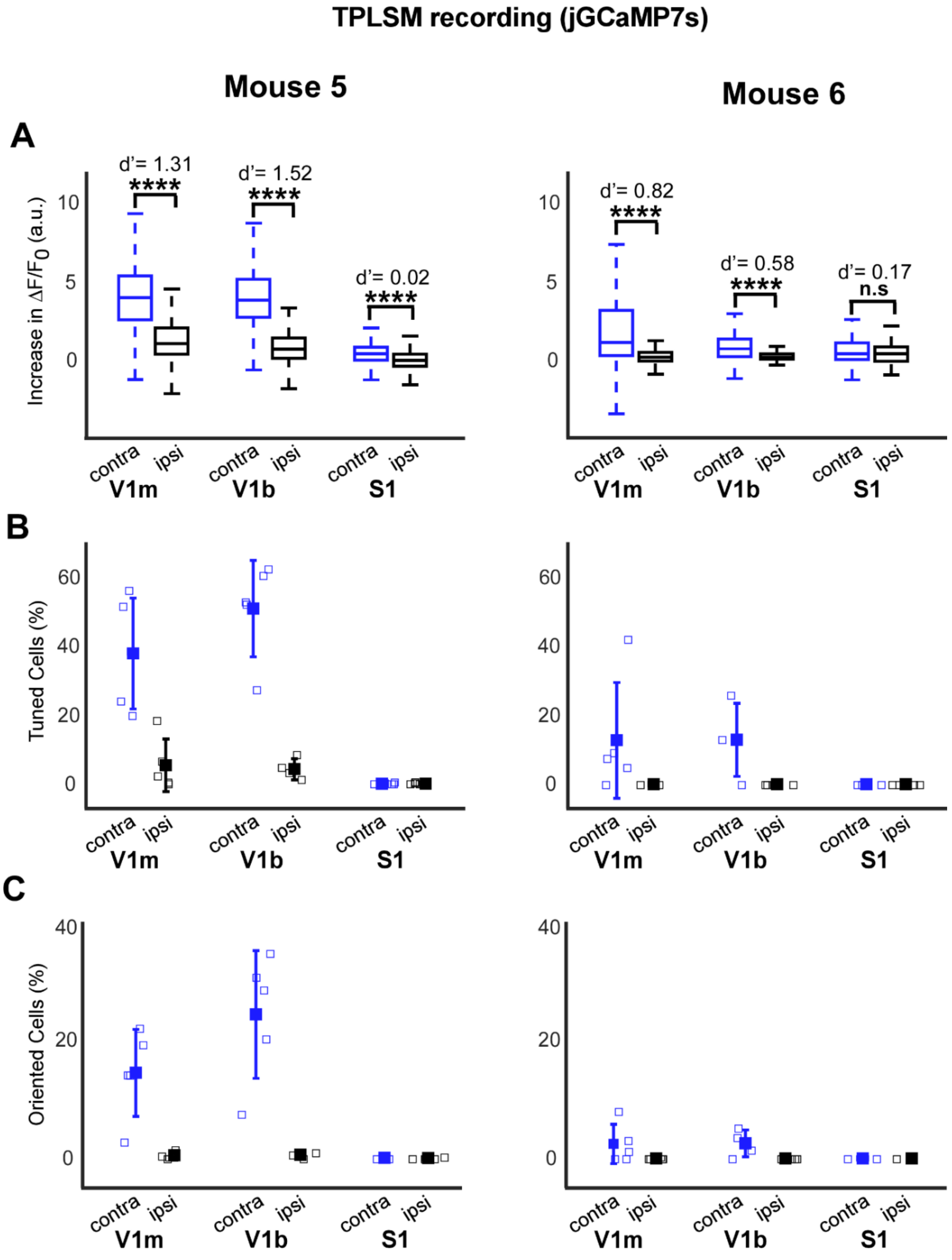


**Supplementary Figure 3: A. Single-mouse data of visual-evoked responses in each brain region showing increased activity in contralateral V1 regions, as recorded with jGCaMP7s.** Recording of increases in jGCaMP7s fluorescence signal during visual stimulation showed similar results to recording with CaMPARI (n=2 mice, see details in Supp. Fig. 3) (Mouse 5: 930-2085 cells/region; median=1433; Mouse 6: 150-339 cells/region, median= 275)

**B.** The fraction of cells with a significant increase in jGCaMP7s fluorescence signal during the presentation of a drifting grating stimulus (tuned cells; ANOVA Test, p=0.01) was higher for contralateral V1 regions (and similar to previously reported data (Dana et al., *Nature Methods*, 2019) than the in the ipsilateral V1 regions. No tuned cells were identified in the S1 regions.

**C.** Cells that showed increased fluorescence changes to different orientations of the drifting grating (oriented cells; ANOVA Test, p=0.01) were found in the contralateral V1 regions.

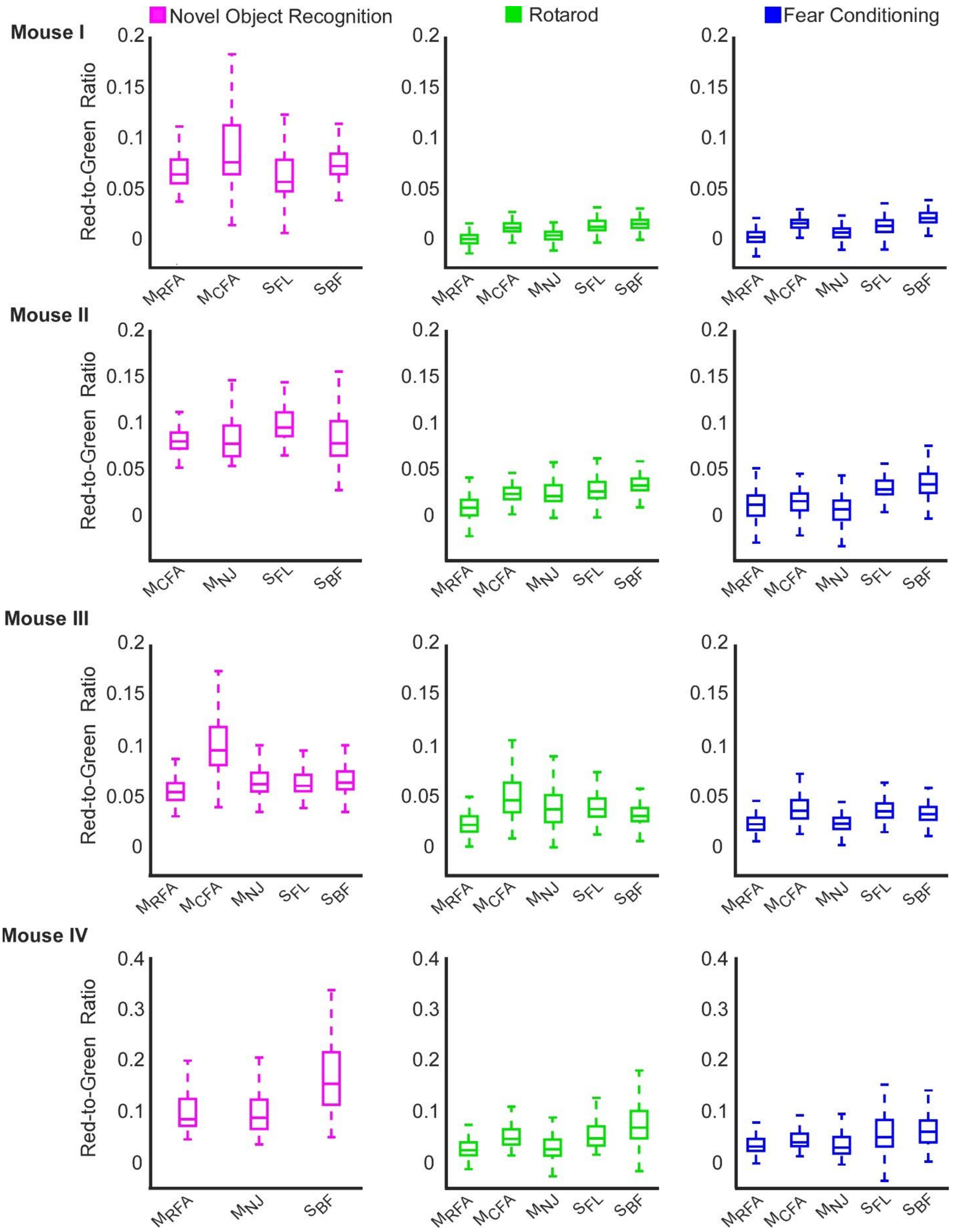

Novel Object Recognition
Rotarod
Fear Conditioning
Mouse I
Mouse II
Mouse III
Mouse IV
Red-to-Green Ratio
MRFA
MCFA
MNJ
SFL
SBF

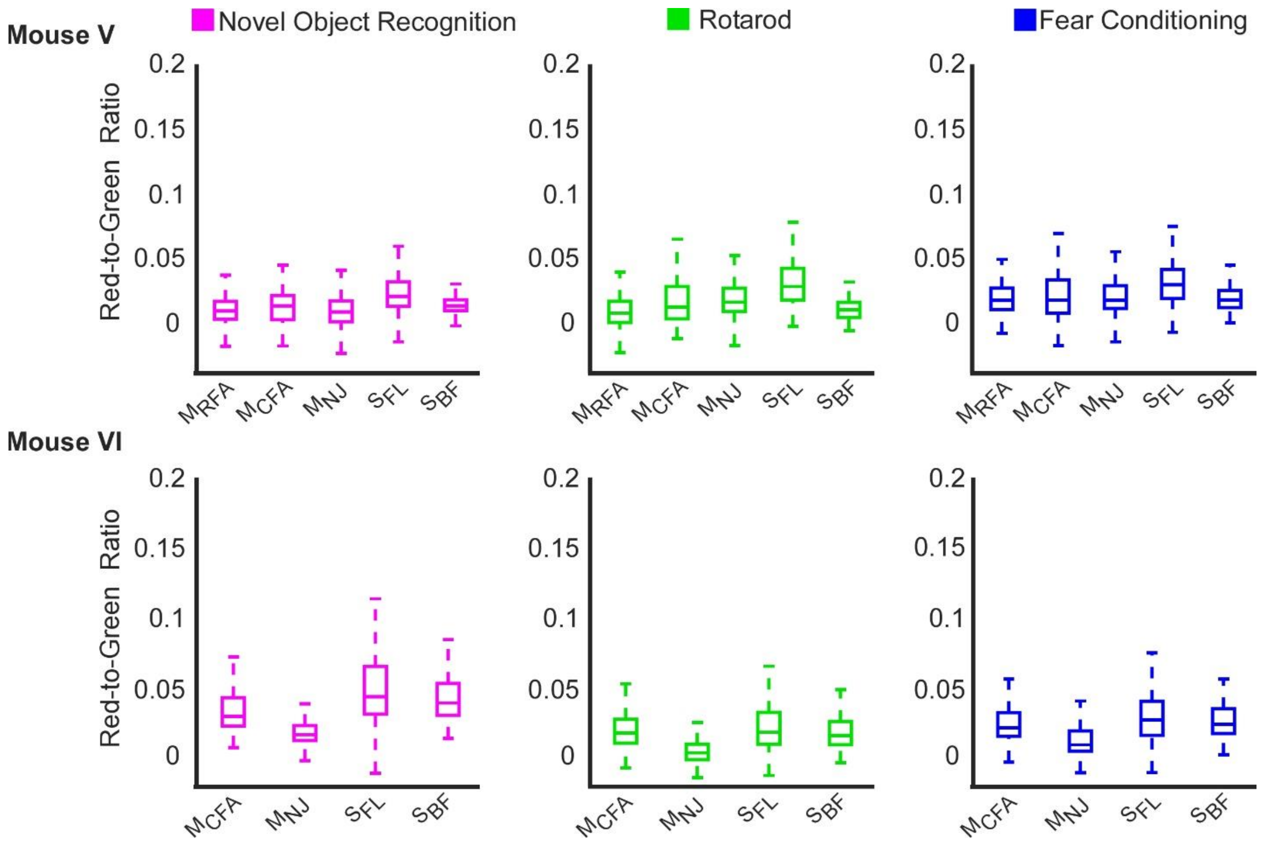


**Supplementary Figure 4: Differential levels of brain activity in individual freely-moving mice during NOR, RR, and FC tasks.** Summary of RGR from all recorded brain regions of all recorded mice (n=6) while performing three different behavioral and cognitive tasks (Mouse I: 95-905 cells/region, median=441; Mouse II:42-763 cells/region, median=144; Mouse III: 166-1315 cells/region, median=597; Mouse IV: 89-559 cells/region, median=177; Mouse V: 94-906 cells/region, median=271; Mouse VI: 116-550 cells/region, median=339). Data from mouse III is presented in Fig. 2D, and the median values from all mice were used in Fig. 2E-G. Note that not all brain regions were recorded in all sessions, since expression levels were either increased with time, or window quality may have deteriorated between recording sessions.

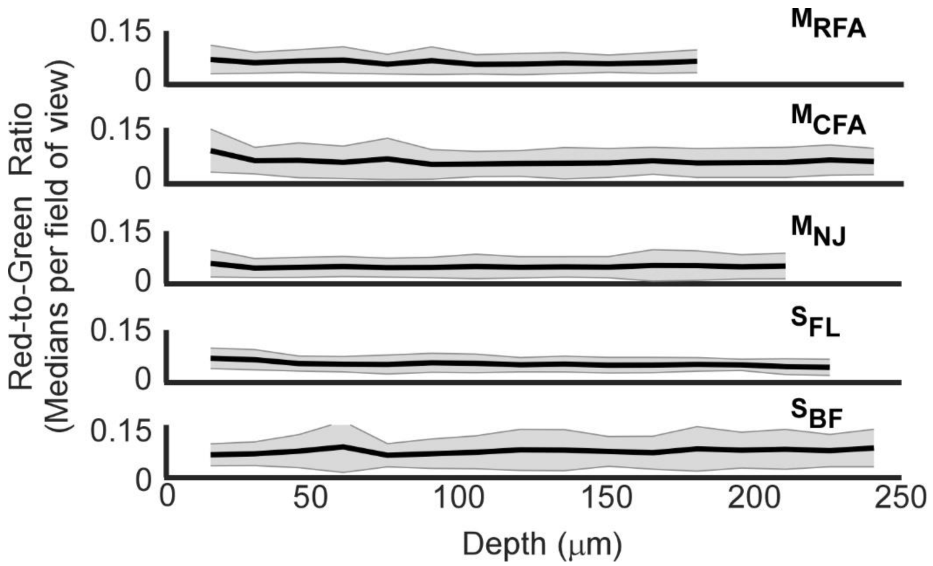


**Supplementary Figure 5: No significant effect of recording depth on RGR levels was identified.** We compared RGR values measured at different tissue depths across different brain areas during the NOR test. RGRs from different brain regions (top panel) at different depth showed no apparent decrease (**$M_{RFA}$:** 4 mice, 5-51 cells/FOV, median number of cells= 17; **$M_{CFA}$:** 3 mice, 1-85 cells/FOV, median= 13; **$M_{NJ}$:** 4 mice, 5-73 cells/FOV, median= 18; **$S_{FL}$:** 3 mice, 4-64 cells/FOV, median= 13; **$S_{BF}$:** 5 mice, 2-110 cells/FOV, median= 23; shaded areas show the standard deviation and solid lines show the average across all median measurement at a specific depth). The bottom panel shows the average of the measured RGRs across all regions down to 240μm under the pia, where all brain regions in which we could measure down to this depth were included in this analysis. The solid line shows the average of median RGRs and error bars show the standard deviation. No significant differences were found between different recording depths (5 mice, 2-110 cells/FOV, median=17; one-way ANOVA, p=1.00)

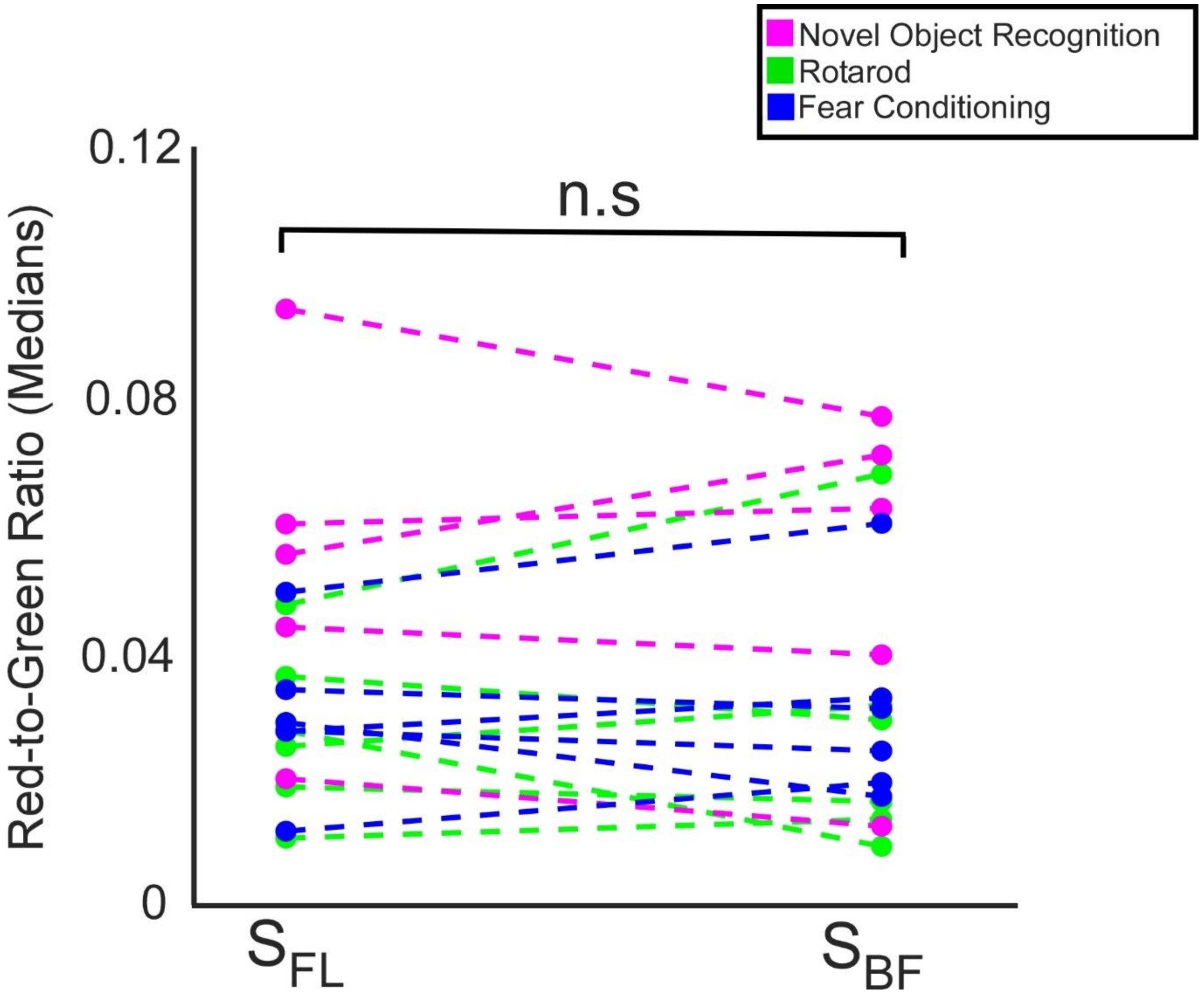


**Supplementary Figure 6: No significant activity changes across the recorded somatosensory regions of freely-moving mice.** Comparison of $S_{FL}$ and $S_{BF}$ during the three tested tasks showed no significant changes (n.s, p=0.96, paired t-test, 42-1315 cells/region, median= 285).

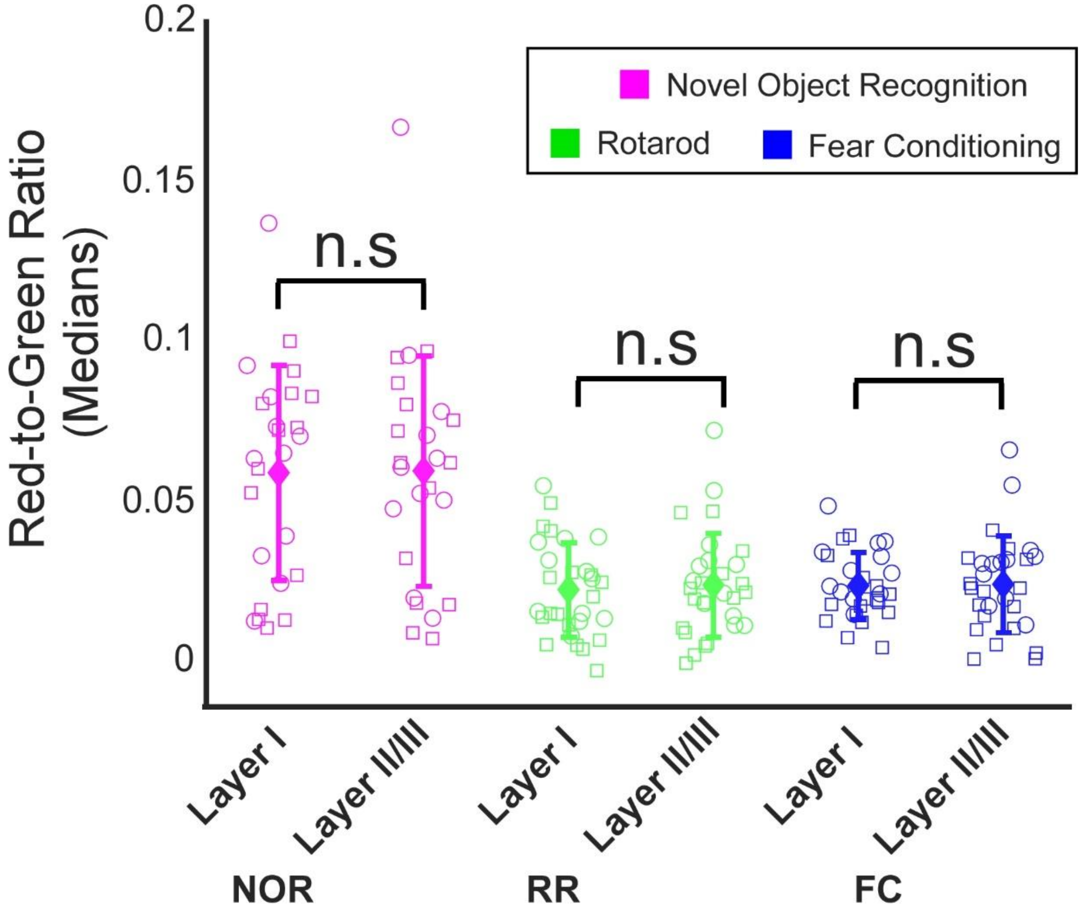


**Supplementary Figure 7: Comparison of activity in layer I and layer II/III cells showed similar activity levels.** We compared the RGR across simultaneously-recorded layer I and layer II/III neurons during NOR, RR, and FC tasks. Activity levels showed similar values and no significant changes were found (**NOR:** Layer I, 9-318 cells/region, median= 77; Layer II/III, 24-765 cells/region, median = 181; p value = 0.79, **RR:** Layer I, 14-183 cells/region, median= 76; Layer II/III, 41-693 cells/region, median = 188; p value = 0.25, **FC:** Layer I, 31-167 cells/region, median= 79; Layer II/III, 47-1189 cells/region, median = 223; p value = 0.76, paired t-tests were used for all comparisons.)

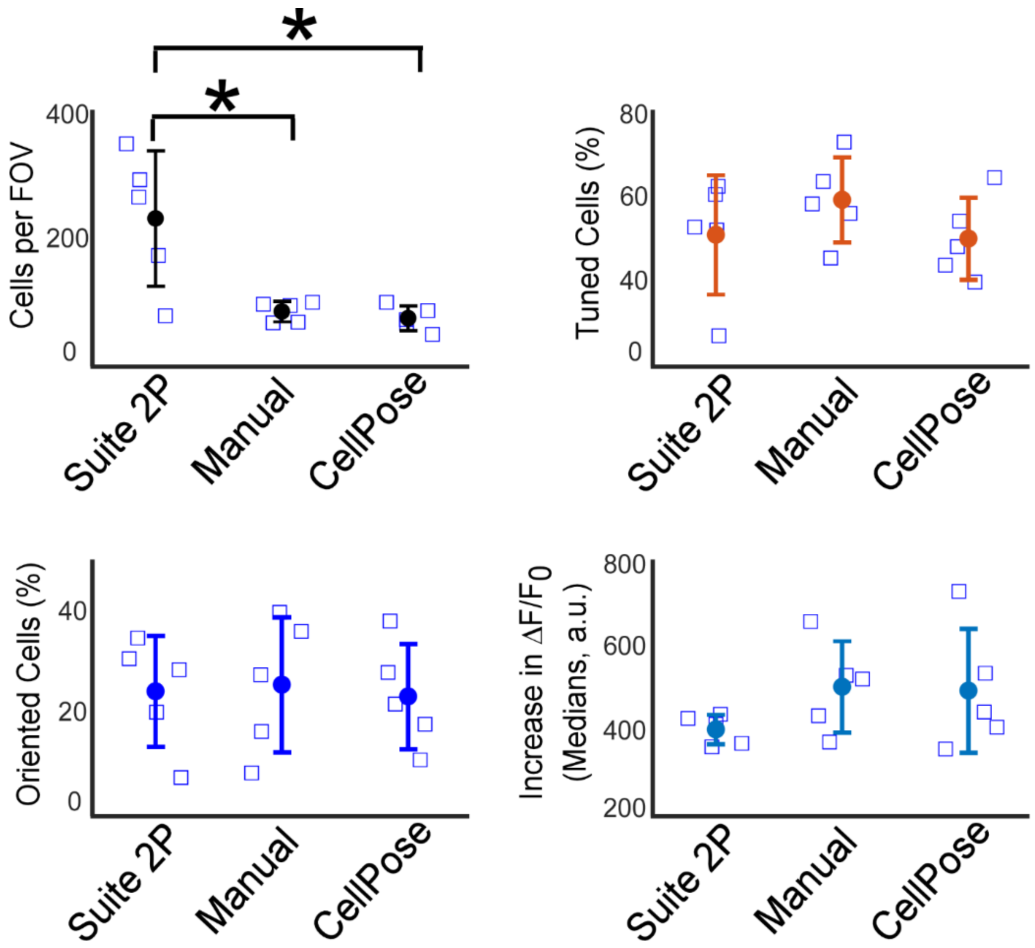


**Supplementary Figure 8: Comparing three cell segmentation methods and their effects on data analysis.** We compared a semi-automated MATLAB script for identifying cells (from Chen et al., *Nature*, 2013) where a user manually identifies cells and the software detects the cell boundaries (manual method) with 2 automated approaches using Suite2P (from Pachitariu et al., *BioRxiv*, 2017), which is designed to extract cell locations from calcium sensors activity movies, and CellPose (Stringer et al., *Nature* Methods, 2021), which is designed to identify cell location from images. All three methods were used to identify cells in the same dataset (n=1 mouse, 5 FOVs) recorded from V1 neurons labeled with jGCaMP7s. Suite 2P detected a significantly larger number of cells from the two other methods (p=0.029 and 0.019 for comparing with manual and CellPose segmentation, respectively; paired t-test), but there were no significant differences in the fractions of tuned or oriented cells, or the identified increases of the fluorescence during visual stimulation.

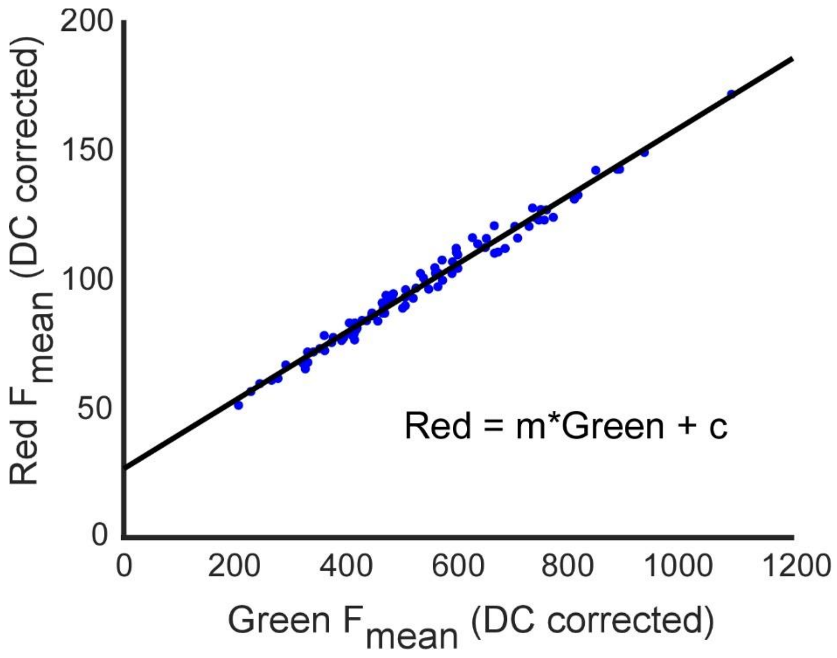


**Supplementary Figure 9: Calculation of RGR values from raw fluorescence data.** Example somatic fluorescence signals from non-photoconverted neurons (after dark current subtraction) showed a linear relationship between the red and green fluorescence signals due to penetration of the green CaMPARI signal into the red channel. To correct for this contamination, we fit the acquired data with a linear function, where the slope (solid black line with slope m and intercept c). The green-to-red contamination factor was calculated by averaging m values from all recordings and was specific for for the filter set that was used to separate the green and red channels. In addition, the line intercept (c) had a small positive value, presumably due to a weak autofluorescence in the red channel and was subtracted as well. m values were calculated separately for the experiments described in Figs. 1 and 2, since they were conducted using different microscopes with different filter sets. For the experiments shown in Fig. 1, c values were calculated for each hemisphere from pre-PC data, and for each recording day. For the experiments described in Fig. 2, we recorded mice before the PC experiments were initiated, and used the average m and c values for the data correction, since no pre-PC data could be recorded on the same day of PC recording.

**Video legends**

**Video 1.** Recording from a freely-moving mouse during the novel object recognition testing. The PC light source was located on top of the enclosed arena and illuminated all of it (the same vertical distance from the arena was maintained across all tasks to provide identical PC conditions), and the mouse was free to move inside it without any mechanical device attached to it.

**Video 2.** Recording from a freely-moving mouse during the rotarod task. Similarly to Video 1, the light source illuminated the entire testing region, and the mouse movement was not restricted within it.

**Video 3.** Recording of a freely-moving mouse during the fear conditioning task. Mice were trained to associate a foot shock with an auditory cue. Activity recording was conducted when the auditory cue was presented to the mouse, and without the foot shock.

**Video 4.** Example data showing *in vivo* volumetric CaMPARI recording with cellular resolution from the surface of the brain down to the bottom of layer II/III (~330 μm). Note that small tissue movements were due to animal breathing during the recording.
